# Sleep controls peroxisomal abundance to reduce wake-induced brain oxidation

**DOI:** 10.1101/2025.11.28.691144

**Authors:** Camilo Guevara, Yongjun Li, Elana Pyfrom, Samantha Killiany, Zhifeng Yue, Amita Sehgal

## Abstract

Sleep is increasingly linked to the regulation of Reactive Oxygen Species (ROS) and lipid metabolism. However, the mechanisms underlying this interaction are underexplored. Here, we use *Drosophila melanogaster* to report a bidirectional relationship between sleep and peroxisomes, cellular organelles that process lipids and alleviate ROS. Of the genes that change expression after sleep deprivation in the dorsal fan-shaped body, knockdown of the peroxisomal biogenesis factor *Pex16* results in decreased sleep. *Pex16* acts in several brain regions to modulate sleep amount, with ellipsoid body neurons (EB) producing the highest sleep reduction of the sleep-promoting regions. Consistent with a general role for peroxisomes, knockdown of other peroxisomal enzymes relevant for lipid import and synthesis also decreases sleep. Whole-brain peroxisomal numbers increase with wake, which is supported by lipidomic analysis indicating that peroxisomal-derived phospholipids are the major contributors to phospholipid changes after wake or sleep deprivation. Peroxisomal proliferation in the EB is driven by neuronal activity and increased oxidation, suggesting that these mediate the effect of wake/sleep loss. In turn, peroxisomes alleviate the oxidation accumulated during wake, such that loss of *Pex16* in the EB works non-cell autonomously to increase lipid peroxidation brain-wide. This likely contributes to sleep loss, as sleep is rescued with an antioxidant. Together, these results position peroxisomes as key players in sleep, regulating ROS and thereby maintaining normal cycles.

## Introduction

Sleep is a universal behavior necessary for survival. However, the cellular and molecular basis of sleep regulation is still debated. Recent studies have suggested a link between sleep, lipid metabolism [1], and ROS regulation [2,3]. At the cellular level, these metabolic processes are performed mainly by the mitochondria, the endoplasmic reticulum (ER), and the peroxisome. While the roles of the mitochondria and the ER in sleep have been explored[1,4,5], the role of peroxisomal physiology in sleep has not been studied.

Peroxisomes are ubiquitous organelles with unique and complementary functions to other lipid and ROS-processing organelles[6–8]. Peroxisomes are involved in the catabolism of branched fatty acids, the synthesis of bile acids, ether lipids, and ROS detoxification; and they collaborate with the mitochondria and ER in the early steps of Very Long Chain Fatty Acids (VLCFA) catabolism and plasmalogen biosynthesis, respectively[9,10]. Peroxisomes are highly important for the nervous system, as illustrated by the strong neurological defects observed in patients with Peroxisomal Biogenesis Disorders (PBDs) and peroxisomal metabolic defects[11,12]. In addition, peroxisomal dysfunction is a common denominator in aging and neurodegenerative diseases[13,14]. Their metabolic functions and their role in brain function position peroxisomes as a strong candidate in sleep regulation.

Here, we use *Drosophila melanogaster* to study the bidirectional relationship between sleep and peroxisomal function. We found that knocking down the peroxisomal biogenesis factor *Pex16* in the dorsal fan-shaped body and in other sleep-regulating regions reduces sleep. Of all the sleep-promoting regions tested, *Pex16* knockdown had the strongest effect in the ellipsoid body (EB); therefore, we targeted the expression of other peroxisome genes to the EB and found that genes necessary for lipid import and ether lipid synthesis are required for sleep. We report that peroxisomal abundance changes throughout the day, with higher abundance after periods of activity or sleep deprivation. The changes in abundance are also reflected in changes in levels of phospholipids, showing that after wake and sleep deprivation, peroxisome-produced phospholipids are the highest contributors to the alteration in neuronal phospholipid species. Wake-dependent increases in peroxisome abundance are likely driven by oxidation derived from neuronal activity, and this is consistent with a role for peroxisomes in ROS detoxification. The loss of *Pex*16 and peroxisomal lipid synthesis in the EB increases brain-wide lipid peroxidation. Antioxidant feeding of *Pex16* knockdowns rescues sleep, indicating that the buildup of oxidative damage is responsible for the sleep loss. These findings demonstrate that during sleep, peroxisomes are important to control wake-dependent ROS accumulation, positioning the peroxisomes as a key player in the interplay between sleep and ROS detoxification.

## Results

### The peroxisomal biogenesis factor 16 (*Pex16*) is relevant for baseline sleep in the dorsal-fan-shaped body

The predominant model for sleep suggests that sleep is regulated by a circadian process, which determines the timing for sleep, and a homeostatic process that accumulates sleep need as a function of time awake[15]. Different brain regions in the *Drosophila* brain play roles in each of these processes, with the dorsal fan-shaped body (dfb) being an important region in the homeostatic process[16,17]. Indeed, twelve hours of mechanical sleep deprivation was shown to produce changes in gene expression in dfb neurons[18]. To test which of these genes are relevant for baseline sleep, we used the R23E10 driver, which includes the dfb, to express RNAi for some of the differentially expressed genes in a developmental and adult-specific manner using the Gal4 (Fig. 1A) and the Gene Switch system (Fig. 1B), respectively. Knockdown of the peroxisomal biogenesis factor 16 (*Pex16*), amnesiac (*amn*), and Eclosion hormone (*Eh*) in males consistently reduced sleep to a similar extent in both systems. Given the increasing evidence linking ROS, lipid metabolism, and sleep, and the known role of peroxisomes in the first two processes[9,10], we decided to focus on the *Pex16* phenotype to explore the role of peroxisomes in baseline sleep regulation. We found that *Pex16* KD reduced sleep in males and females (Fig. 1C-F, S1 Fig A-B).

**Figure 1:**
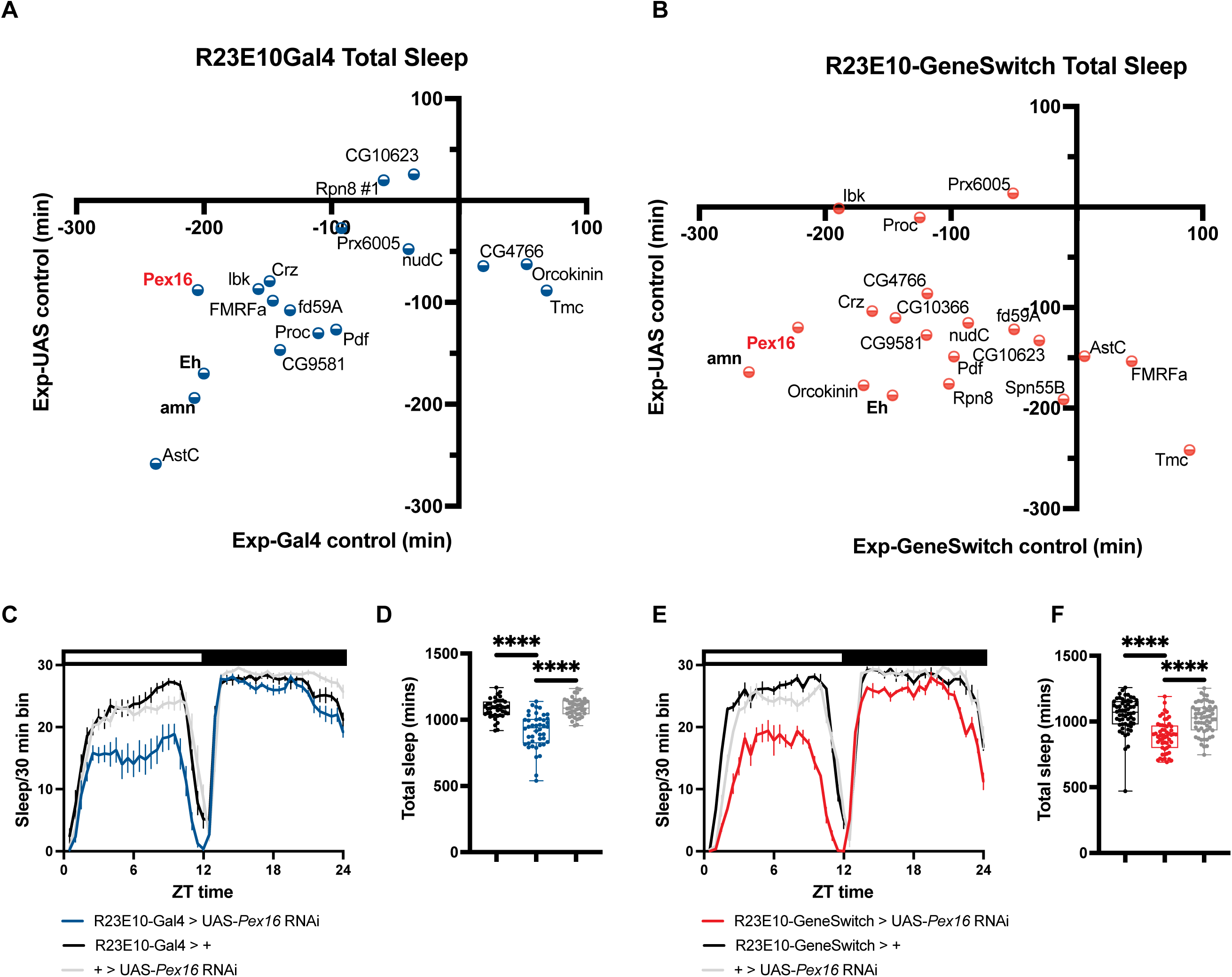
Developmental and adult-specific knockdown of Pex16 with the R23E10 driver reduces sleep. **(A)** Sleep differences after knockdown of sleep deprivation differentially expressed genes in the dorsal fan-shaped body with the R23E10Gal4 driver (n=14-16). (**B)** Sleep differences after knockdown of the same genes in (A) in the R23E10GeneSwitch driver (n=14-16). X and Y axes for A and B represent the sleep differences between the knockdown group and the Gal4/GeneSwitch and UAS control, respectively. (**C)** Male sleep trace for Pex16 knockdown using the R23E10Gal4 driver. (**D)** Total sleep time after Pex16 knockdown in the R23E10Gal4 driver (n = 42-43). ****p < 0.0001 by Ordinary one-way ANOVA with Bonferroni multiple comparisons correction. (**E)** Male sleep trace for Pex16 knockdown in the R23E10GeneSwitch driver. (**F)** Total sleep time after Pex16 knockdown in the R23E10GeneSwitch driver (n = 57-58). ****p < 0.0001 by Kruskal-Wallis test with Dunn’s multiple comparisons correction.

### Knockdown of peroxisomal biogenesis reduces sleep in sleep-relevant neuronal types

Given that peroxisomes are ubiquitous organelles highly important for neuronal function[12], and recent reports show that the sleep effects of the R23E10 driver also originate from the Ventral Nerve Cord (VNC),[19,20] we performed *Pex16* knockdown in different brain cells, including sleep-regulating neuronal subpopulations and glia. We found that *Pex16* knockdown in sleep-promoting mushroom body output neurons (MB05B1-Gal4)[21] and ellipsoid body neurons (R58H05-Gal4)[22] reduced sleep. It also did so in wake-promoting mushroom body output neurons (MB011B-Gal4)[21] and dopaminergic neurons (c584-Gal4)[23]. Interestingly, pan-neuronal and pan-glial knockdown of *Pex16* resulted in small decreases or no changes in sleep (Fig. 2A). Consistent with a potential role in sleep-regulating circuits, *Pex16* knockdown in Hugin neurons, which are not sleep nor wake-promoting[24], resulted in a small sleep reduction in males and no changes in sleep in females (Fig. 2A).

**Figure 2:**
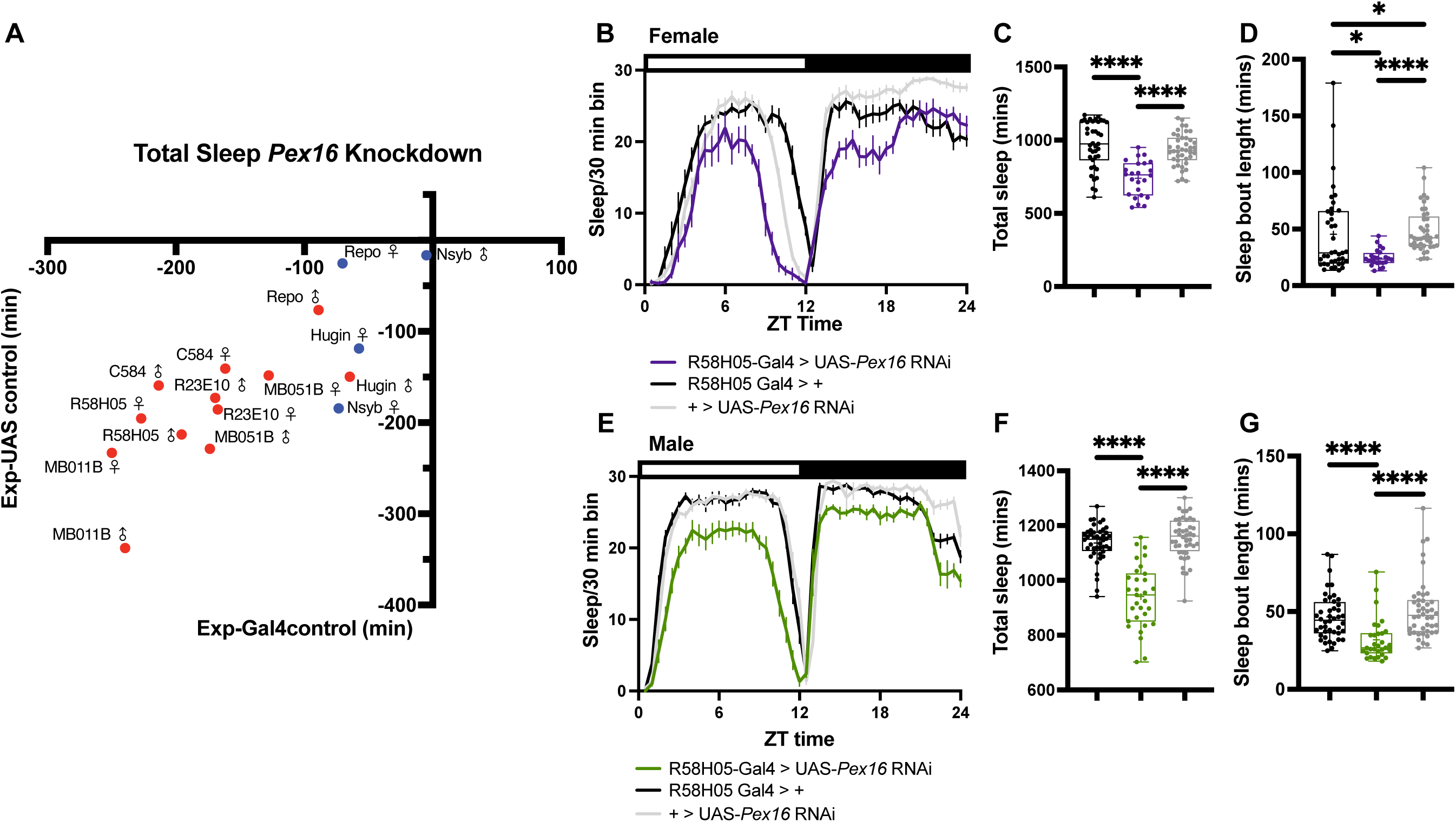
Pex16 knockdown in sleep-relevant regions reduces sleep, with strong effects in the ellipsoid body. **(A)** Sleep differences after knockdown of Pex16 in different brain cell populations in males and females. X and Y axes represent the sleep differences between the knockdown group and the Gal4 and UAS control, respectively. Red dots indicate significant sleep time differences compared with the UAS and Gal4 controls (n=31-60), p < 0.05 by One-way ANOVA with Bonferroni multiple comparisons correction or Mann-Whitney Test with Dunn’s multiple comparisons correction, while blue dots represent non-significant differences. R23E10: dorsal fan-shaped body and ascending neurons driver; R58H05: ellipsoid body driver; MB051B: Sleep-promoting mushroom body output neurons driver; MB011B: Wake-promoting mushroom body output neuron; Hugin: hugin neurons driver; Repo: pan-glial driver; c584: dopaminergic neurons. (**B)** Female sleep trace for Pex16 knockdown in the R58H05 driver. **(C-D)** Sleep parameters affected by Pex16 knockdown in the R58H05 driver in females, total sleep time (C) (n= 39-41), and average sleep bout length (D) (n= 39-41). **E)** Male sleep trace for Pex16 knockdown in the R58H05 driver. **(F-G)** Sleep parameters affected by Pex16 knockdown in the R58H05 driver in males, total sleep time (F) (n= 31-44), and average sleep bout length (G) (n= 31-44). For figures (C-D, E-F) *p < 0.05, ****p < 0.0001 by Kruskal-Wallis test with Dunn’s multiple comparison correction.

Given the known role of ellipsoid body (EB) neurons in sleep regulation[22,25,26] and their physiological response to extended wake[22,26], together with the dramatic sleep loss produced after *Pex16* knockdown in these neurons, we decided to focus on the effect of peroxisomal dysfunction in the EB. The Brain Single-Cell Transcriptome atlas[27] data show that the gene corresponding to the promoter used to target these neurons (*CG6024*), as well as *Gad1,* which is expressed in EB neurons,[28] colocalize with *Pex16* and the peroxisomal lipid transporter *Pmp70* (S2 Fig A). We found that *Pex16* knockdown in the EB resulted in a decrease in total sleep and a reduction in sleep bout length (Fig. 2B-G). To assess the efficiency of *Pex16* RNAi, we used a GFP with a peroxisomal localizing signal (GFP-SKL), which forms puncta within peroxisomes when they are present[29]. When GFP-SKL is expressed in the ellipsoid body neurons, it is distributed in punctate structures in the cell body and the ring (S2 Fig B), indicating GFP accumulation in peroxisomes. On the other hand, when we co-expressed *Pex16* RNAi and GFP-SKL, we saw a diffuse GFP signal with no evidence of punctate structures in the ring or cell bodies (S2 Fig B), suggesting the absence of peroxisomes. This is consistent with the role of *Pex16* in peroxisomal biogenesis and confirms the efficiency of the RNAi used.

To verify that the effects observed were from the EB, we inhibited potential VNC contribution using *teashirt-*Gal80 and found that brain-only activation of the EB neurons using the R58H05 driver increased sleep (S2 Fig C-D). We also performed a *Pex16* knockdown using a split-Gal4 line that is expressed only in the EB within the brain (S2 Fig E), and this reduced sleep consistent with the broader EB driver (S2 Fig F-G). Finally, we counted cell numbers after *Pex16* knockdown and found no differences between the knockdown and control groups (S2 Fig H-I), suggesting that peroxisomal knockdown does not compromise cell viability.

### Peroxisomal lipid import and ether-synthesis modulate baseline sleep

Lipid metabolism is a promising link between peroxisomal function and sleep. As we found that peroxisomal biogenesis is essential for sleep, we asked what metabolic pathways involved in peroxisomal lipid metabolism modulate baseline sleep. Thus, we knocked down the long-chain fatty acid transporter *Abcd3* (ATP-binding cassette subfamily D member 3), the very long-chain fatty Acid (VLCFA) β-oxidation enzyme *Acox1* (acyl-CoA oxidase 1), and the ether-lipid synthesis enzyme *Gnpat* (Glycerophosphate O-acyltransferase) in the ellipsoid body neurons. Both *Abcd3* and *Gnpat* knockdown reduced sleep in males and females (Fig. 3 A-H), but *Acox1* only reduced sleep significantly in males (Fig. 3 I-L). When the same genes were knocked down using additional sleep-promoting drivers, smaller or non-significant effects were observed after *Acox1* knockdown, while *Abcd3* and *Gnpat* knockdown consistently reduced sleep (S3 Fig A-C). These results suggest that peroxisomal lipid synthesis and import are necessary for normal baseline sleep levels.

**Figure 3:**
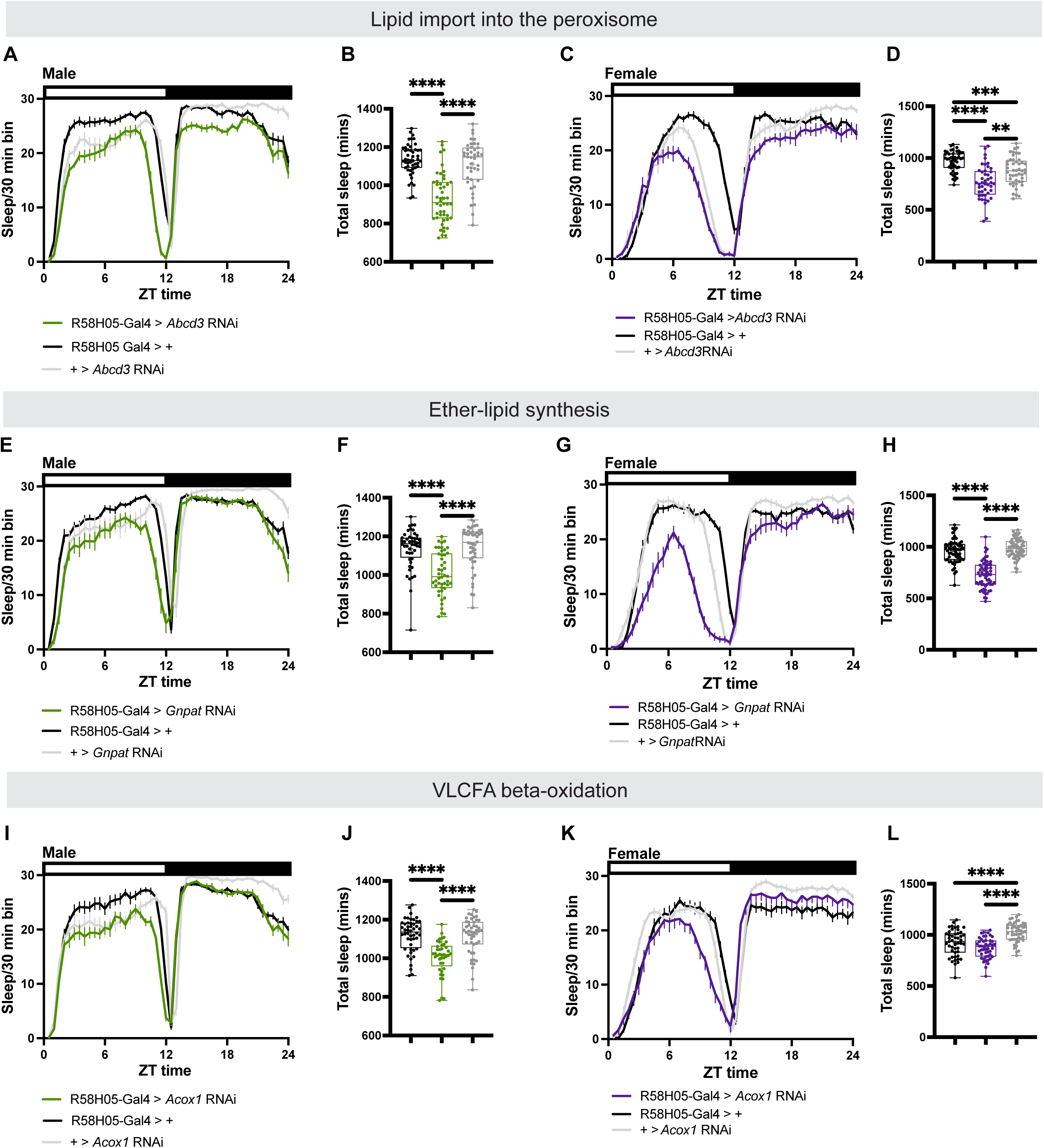
Knockdown of peroxisomal lipid import and synthesis, but not beta-oxidation, in the ellipsoid body reduces sleep. **(A, E, I)** Male sleep traces after knockdown of Abcd3 (A), Gnpat (E), and Acox1 (I) in the R58H05 driver. **(B, F, J)** Total sleep time after knockdown of Abcd3 (B) (n= 43-48), Gnpat (F) (n= 46-48), and Acox1 (J) (n= 41-48) in males. **(C, G, K)** Female sleep traces after knockdown of Abcd3 (C), Gnpat (G), and Acox1 (K) in the R58H05 driver. **(D, H, L)** Total sleep time after knockdown of Abcd3 (D) (n= 46-47), Gnpat (H) (n= 61-64), and Acox1 (L) (n= 47-48) in females. For figures (B, D, H), **p < 0.01, ****p < 0.0001 by Brown-Forsythe ANOVA test with Dunnett’s multiple comparison correction. For figures (F, J) ****p < 0.0001 by Kruskal-Wallis test with Dunn’s multiple comparison correction. For Figure (L) ****p < 0.0001 by Ordinary one-way ANOVA with Bonferroni multiple comparisons correction in females.

### Peroxisome abundance changes throughout the day and is modulated by sleep

Finding that peroxisomal metabolism, particularly lipid synthesis, is important for baseline sleep, we sought to determine how sleep modulates peroxisomal abundance and metabolism. *Pex16* is one of the main peroxisomal biogenesis factors, and it is relevant for the insertion of proteins in the peroxisomal membrane[9]. The lack of *Pex16* results in the loss of or non-functional peroxisomes in different eukaryotic models[29–31] and humans[32]. Given the changes in *Pex16* expression after sleep deprivation[18] and its effects on baseline sleep, we evaluated how peroxisomal abundance changes throughout the day and whether it is modulated by sleep deprivation. To test this, we used a cerulean-fluorescently tagged peroxisomal membrane protein (PMP34-mCerulean), which has been used previously as a peroxisomal marker[33], to measure brain-wide peroxisomal abundance after the morning and evening locomotor activity peaks (ZT2 and ZT14), and after daytime siesta and nighttime sleep (ZT8 and ZT20) (S4 Fig A-B). We found that peroxisomal abundance is highest after wake periods and is reduced after sleep (Fig. 4A-B, S4 Fig C). This result suggests that peroxisomal abundance increases as a function of activity and that after sleep, peroxisomal numbers decrease.

**Figure 4:**
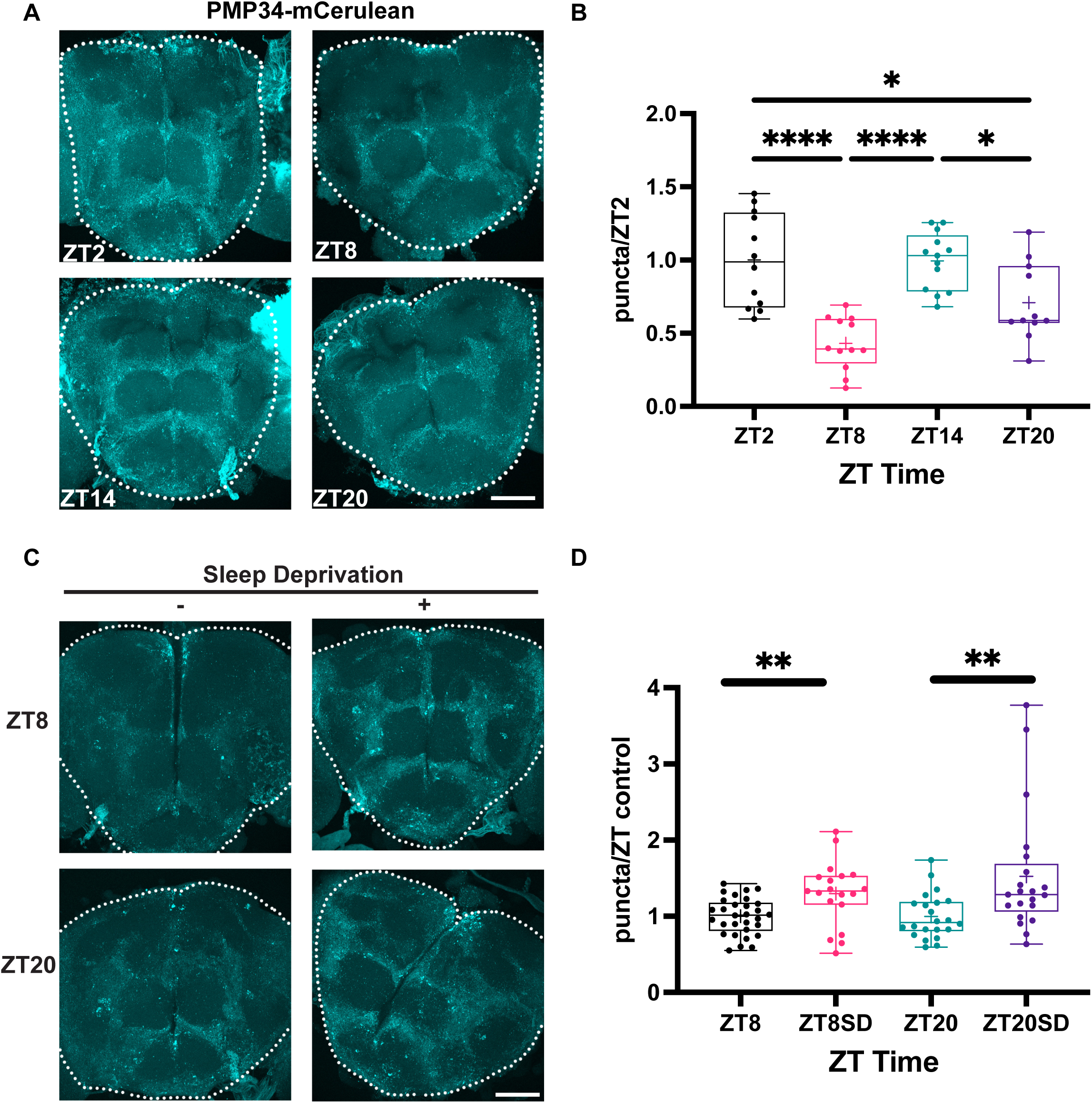
Peroxisomal abundance changes during the day and increases after sleep deprivation. **(A)** Representative images of maximum intensity projections of PMP34-mCerulean brains at the given timepoints; the dotted area corresponds to the region analyzed. **(B)** Quantification of mCerulean puncta relative to ZT2 (n= 11-13), *p < 0.05, ****p < 0.0001 by Ordinary one-way ANOVA with Bonferroni multiple comparisons correction. **(C)** Representative images of maximum intensity projections of PMP34-mCerulean brains at the given points with (+) and without (-) six hours of mechanical sleep deprivation; the dotted area corresponds to the region analyzed. **(D)** Quantification of mCerulean puncta relative to their corresponding non-sleep-deprived control (n= 19-30), **p < 0.01 by unpaired t-test between ZT8 and ZT8SD, **p < 0.01 by Mann-Whitney test between ZT20 and ZT20SD. Scale bar is 50μm.

To further evaluate the modulation of peroxisomal abundance with sleep, we mechanically sleep-deprived flies for six-hour periods, between ZT2 and ZT8, and between ZT14 and ZT20, to prevent the daytime siesta and nighttime sleep episodes, respectively. We found that mechanical sleep deprivation increases peroxisomal numbers in males (Fig. 4C-D). Surprisingly, we found no effect in females (S4 Fig D). These results demonstrate that peroxisomal abundance changes throughout the day in a sleep-dependent fashion, and suggest that male flies, in particular, rely on peroxisome function after extended wake.

### Phospholipid changes after extended wake reflect peroxisomal involvement

Peroxisomes are highly metabolic organelles involved in lipid metabolism[9,10]. And we demonstrated that lipid synthesis is an important peroxisomal metabolic process for baseline sleep regulation. Therefore, changes in peroxisomal gene expression and abundance after wake and sleep deprivation should be reflected in changes in peroxisome-related lipid composition. Lipidomic analysis of sorted neurons and glia reveals that sleep need influences the abundance of phospholipids, diacylglycerols (DAGs), and triacylglycerols (TAGs) in both neurons and glia[34]. Interestingly, most phosphatidylcholine (PC) and Phosphatidylserine (PS) species that changed in abundance as a function of sleep need were detected in neurons[34]. As *Pex16* knockdown in various neuronal subpopulations reduced sleep, with no effects in glia, we focused largely on peroxisomal phospholipid abundance in neurons after wake and sleep deprivation.

Peroxisomes are responsible for the synthesis of ether-lipids and are necessary for the early steps of plasmalogen biosynthesis[9,10]. Therefore, we examined the proportion of phospholipids that changed abundance relative to ZT2 after wake (ZT14) or after sleep deprivation (ZT2SD) and contained either a plasmalogen or an ether lipid in any of their fatty acid side chains. We used an FDR<0.01 as our inclusion criteria. Interestingly, after a day of wake (ZT14), ∼42% and 95% of the significant PC species that changed in abundance relative to ZT2 were peroxisome-generated in females and males, respectively (Fig. 5A), while all the significant PS species presented an ether lipid or plasmalogen-containing tail (Fig. 5B). In sleep-deprived conditions, all the neuronal phospholipid species with significant abundance changes relative to ZT2 were peroxisome-generated (Fig. 5C). Remarkably, and consistent with the increase in peroxisomal abundance after wake or sleep deprivation, both PC and PS, except for PS(O-29:0), increased their abundance after those conditions. In female glia at ZT14, 75% of the phosphatidylethanolamines (PE) that changed abundance relative to ZT2 were peroxisome-related (S5 Fig A). After sleep deprivation, only one significant PE species was detected in female glia, and this is potentially peroxisome-related (PE (37:7), PE(P-38:6), PE (36:0), PE(O-37:0)). However, for male glia, sleep state does not affect phospholipid abundance. The changes in peroxisomal gene expression, abundance, and peroxisomal-related lipids after extended wakefulness suggest a modulatory role of sleep in determining peroxisomal abundance and metabolism.

**Figure 5:**
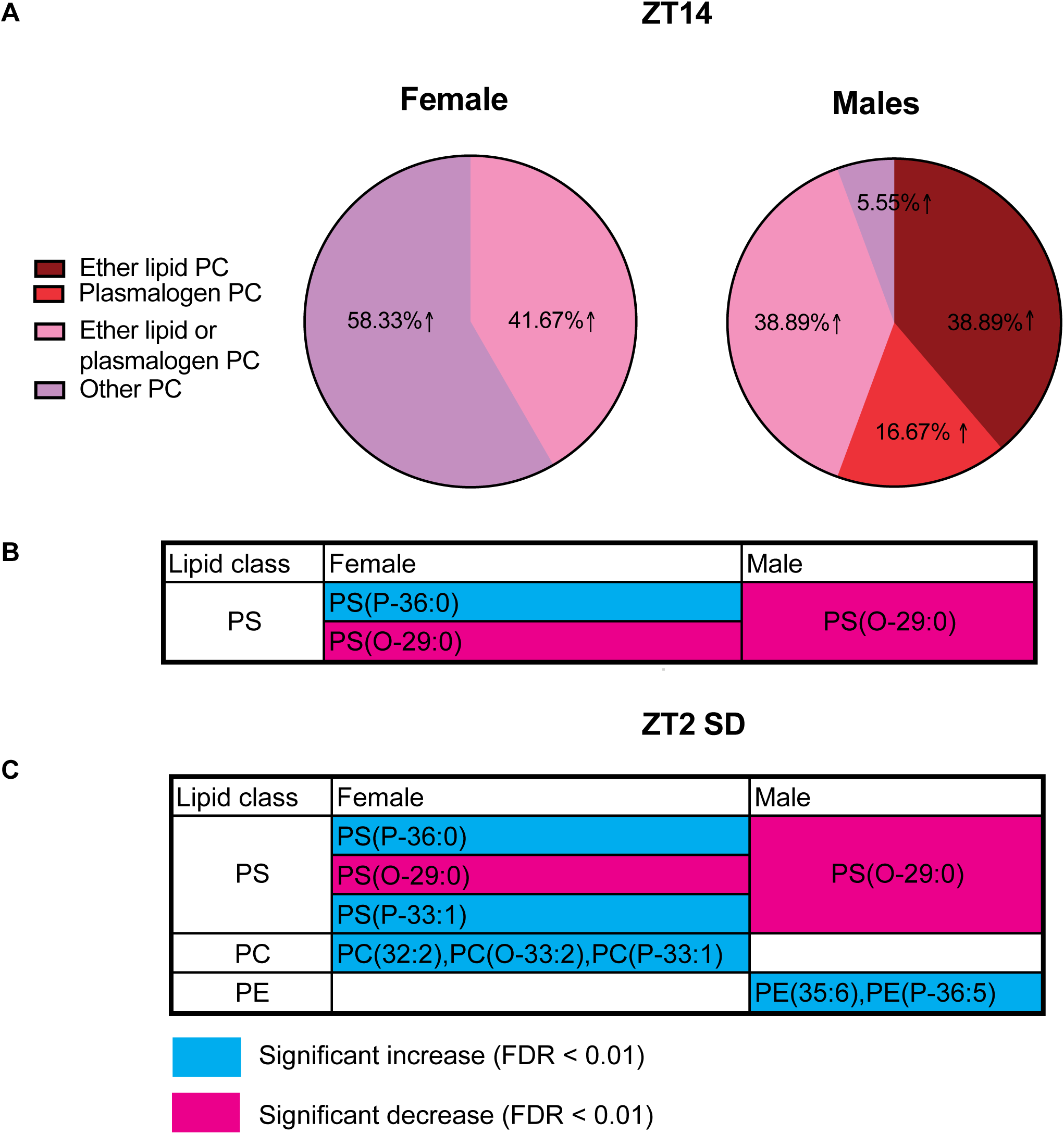
Peroxisomal-derived phospholipids change their abundance after extended wake in neurons. (A) Percentage of peroxisomal-related phosphatidylcholines (PC) showing a significant change in abundance compared to ZT2 in neurons. The arrow shows an increased abundance relative to ZT2. (B) phosphatidylserines (PS), showing a significant change in abundance compared to ZT2 in neurons. **(C)** Phospholipids exhibit a significant change in abundance in neurons after 8 hours of mechanical sleep deprivation. PE: phosphatidylethanolamine. P-: plasmalogen, O-: ether-linked lipid

### Neuronal activity and oxidation increase peroxisomal abundance in the ellipsoid body

To explore how extended wake affects peroxisomal proliferation, we sought to evaluate peroxisomal dynamics after physiological manipulations of the ellipsoid body. After sleep deprivation, there is an increase in neuronal activity of the EB and in the expression of synaptic markers in its neuropil ring[22]. Therefore, we asked whether the activity of the ellipsoid body was relevant for peroxisomal proliferation. We activated the ellipsoid body neurons using TRPA1 and measured changes in GFP-SKL as a proxy for peroxisomal abundance. We found that after activation, which increased sleep as expected (S6 Fig A), there was an increase in GFP intensity in the ring (Fig. 6A-B), suggesting an increase in peroxisomal abundance. Notably, the increased sleep does not decrease peroxisomes as consistent activation of the EB maintains pressure for sleep.

**Figure 6:**
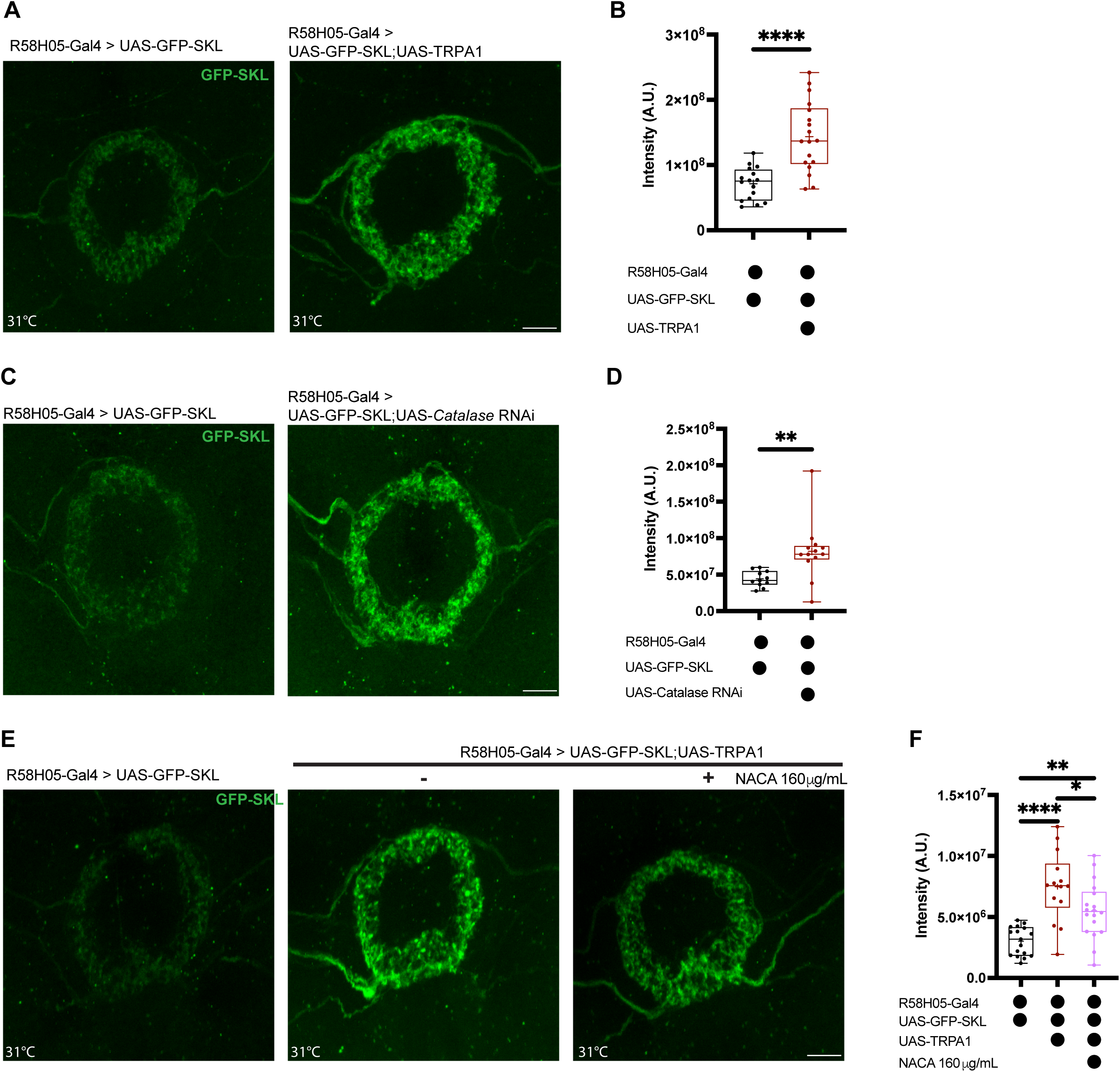
Activity-derived oxidation induces peroxisome proliferation. **(A)** Representative images of maximum intensity projections of the ellipsoid body ring neuropil expressing peroxisomal GFP (GFP-SKL) after 24-hour exposure to 31°C with (right) and without (left) thermogenetic ellipsoid body activation. **(B)** GFP-SKL intensity quantification in the ring neuropil with the conditions described in (A) (n= 16-18), ****p < 0.0001 by unpaired t-test. **(C)** Representative images of maximum intensity projections of the ellipsoid body ring neuropil expressing GFP-SKL with (right) and without (left) catalase knockdown. **(D)** GFP-SKL intensity quantification in the ring neuropil with the conditions described in (C) (n= 11-13), **p < 0.01 by Mann-Whitney test. **(E)** Representative images of maximum intensity projections of the ellipsoid body ring neuropil expressing GFP-SKL after 24-hour exposure to 31°C with (center, right) and without (left) thermogenetic ellipsoid body activation. The groups with thermogenetic activation of the ellipsoid body were fed with complete food (center) or complete food supplemented with 160 μg/mL NACA (right). **(F)** GFP-SKL intensity quantification in the ring neuropil with the conditions described in (E) (n= 14-18), *p < 0.05, **p < 0.01, ****p < 0.0001 by Ordinary one-way ANOVA with Bonferroni multiple comparisons correction. Scale bar is 10μm.

Neuronal activity increases oxidation, and oxidation is a trigger for peroxisomal biogenesis[35,36], making oxidation a candidate for linking neuronal activity during wake[18,22,37] to peroxisome proliferation. To test how oxidation affects peroxisomal biogenesis in the ellipsoid body, we knocked down catalase, which increased GFP-SKL intensity in the ellipsoid body ring (Fig. 6C-D). We then asked if neural activation promotes peroxisomal proliferation via oxidation. To test this, we drove expression of the TRPA1 channel in the EB and measured GFP-SKL intensity with and without supplementation of the blood-brain-barrier-permeable[38,39] antioxidant N-acetylcysteine amide (NACA). As previously shown, EB activation with TRPA1 increased GFP-SKL intensity (Fig. 6E-F). And consistent with an interplay between neuronal activity and oxidation, NACA feeding reduced GFP-SKL intensity after EB activation, although it was still higher than that of the non-activated control (Fig. 6E-F). These results, together with the peroxisomal increases observed after behavioral activity and sleep deprivation, suggest that peroxisomes are upregulated in conditions of high neuronal activity and oxidative stress, potentially helping to alleviate the oxidative stress accumulated during wakefulness.

### Peroxisomes regulate brain oxidation and sleep

Once a relationship between extended wake, peroxisomal abundance, and oxidation was established, we asked how brain oxidation was affected in the context of *Pex16* knockdown. To test this, we measured levels of the lipid peroxidation product malondialdehyde (MDA), which have been previously shown to increase after wakefulness[1] and serve as a measurement of oxidative stress[40]. Surprisingly, we found that *Pex16* knockdown in the ellipsoid body increased whole-brain lipid peroxidation in females (Fig. 7A-B) and in males (S7 Fig A-B). As an additional method to measure lipid peroxidation, we used the Bodipy^TM^ C9/C11 dye and found that loss of *Pex16* in the EB increased the ratio between peroxidized and neutral lipids, indicating an overall increase in lipid peroxidation (S7 Fig C-D). MDA levels also increased after *Gnpat* knockdown with respect to the Gal4 control (Fig. 7C-D). Importantly, the increase in MDA levels is not a consequence of sleep loss because feeding the flies with the sleep-promoting GABA agonist Gaboxadol did not reduce lipid peroxidation (S7 Fig E-F), and *Abcd3* knockdown, which reduces sleep as well, resulted in no changes in MDA levels (Fig. S7 Fig G). These results suggest that sleep loss is not the determinant of lipid peroxidation increases, and ether-lipid synthesis is the most important metabolic process in the peroxisome that controls oxidative damage and sleep.

**Figure 7:**
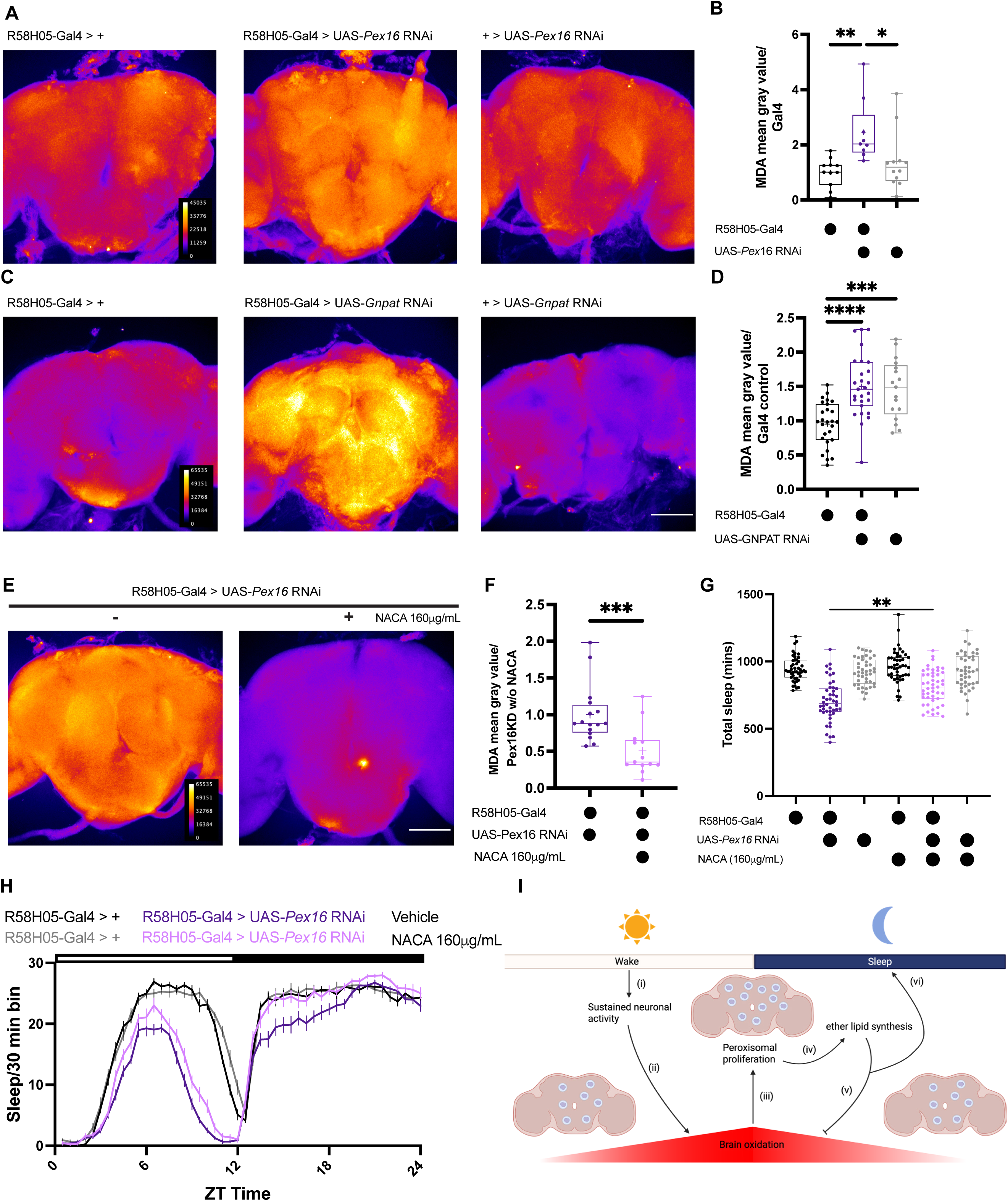
Peroxisomes regulate normal sleep/wake cycles through modulation of brain oxidation. **(A)** Representative images of the sum of intensity projections of brains immunostained for MDA with (center) and without (left, right) Pex16 knockdown with the R58H05 driver. **(B)** Quantification of the mean gray value normalized to the Gal4 control in the conditions described in (A) (n = 14-22), **p < 0.01 by Kruskal-Wallis test with Dunn’s multiple comparisons test. **(C)** Representative images of the sum of intensity projections of brains immunostained for MDA with (center) and without (left, right) Gnpat knockdown with the R58H05 driver. **(D)** Quantification of the mean gray value normalized to the Gal4 control in the conditions described in (C) (n= 17-27), ***p < 0.001, ****p < 0.0001 by Ordinary one-way ANOVA with Bonferroni multiple comparisons correction. **(E)** Representative images of the sum of intensity projections of Pex16 knockdown brains immunostained for MDA with (+) and without (-) NACA 160 μg/mL supplementation in complete food. **(F)** Quantification of the mean gray value normalized to the non NACA fed group (n= 13-16), ***p < 0.001 by Mann-Whitney test. **(G)** Total sleep time after Pex16 knockdown in the R58H05 driver and NACA 160 μg/mL supplementation in complete food (n= 42-47), **p < 0.01 by Brown-Forsythe ANOVA test with Dunnett’s multiple comparison correction. **(H)** Female sleep trace for Pex16 knockdown in the R58H05 driver with and without NACA 160 μg/mL supplementation in complete food. (**I)** A model describing how wake (i), through neuronal activity-dependent oxidation (ii), induces peroxisomal proliferation (iii) and ether lipid synthesis (iv), which potentially reduces brain oxidation (v) and helps maintain normal sleep/wake cycles (vi). For figures (A, C, E), intensity is decoded with a fire LUT, and the scale bar is 100 μm.

To determine whether the peroxisomal regulation of ROS modulates baseline sleep, we fed flies NACA while knocking down *Pex16* expression. NACA feeding reduces whole-brain MDA levels in flies with low Pex16 expression (Fig. 7E-F) and partially rescues total sleep in females, with higher effects for nighttime sleep, and no significant effects in the control groups (Fig. 7G-H) or in male flies (S7 Fig H-I). Altogether, these results suggest that during sleep, peroxisomes are necessary to alleviate the oxidative damage accumulated after wake.

## Discussion

The relationship between sleep and oxidative stress is an area of considerable interest. Sleep deprivation results in oxidative damage in the gut[3], sleep mutants are more susceptible to oxidative stress[2], and after a day of wake, the brain accumulates oxidative damage in the form of peroxidized lipids[1]. In fact, lipids are a known cellular substrate for ROS. Prolonged neuronal activity induces lipid peroxidation of fatty acids *in vitro*[41] and *in vivo*[42], and lipid peroxidation products accumulate after sleep deprivation and regulate the physiology of sleep-promoting neurons[5]. As the primary biochemical functions of the peroxisome are lipid metabolism and ROS detoxification, peroxisomes are prime candidates for having a role in sleep regulation. Indeed, we found that sleep modulates peroxisomal abundance and metabolism, and peroxisomal lipid synthesis in neurons regulates sleep. We propose that wake and sleep deprivation induce peroxisomal proliferation (Fig. 4), primarily through sustained neuronal activity and an increase in oxidation (Fig. 6). This increase in abundance is accompanied by changes in ether-lipid and plasmalogen synthesis (Fig. 5), which modulate sleep (Fig. 3). We propose that during sleep, peroxisome metabolism helps to reduce the brain oxidative damage, and such regulation is necessary to maintain normal sleep/wake cycles (Fig. 7). Neuronal peroxisomes have previously not been implicated in sleep (Fig. 7G).

The sleep reduction observed after loss of peroxisomal genes is consistent with clinical and self-reported studies that report sleep disturbances in patients with peroxisomal syndromes[43,44]. Also, metabolic studies in humans and animal models show changes in peroxisomal metabolites after sleep deprivation[45,46], and ether lipids are one type of lipid that has been associated with sleep disturbances[47]. This work explains these findings from a mechanistic perspective, showing that peroxisomal metabolism and sleep have a bidirectional regulation mediated by the regulation of wake-induced brain oxidation.

Our results are consistent with previous studies, which show that wake and sleep loss induce oxidation and that one of the functions of sleep is to reduce this oxidative burden[1,2,5]. We found that loss of peroxisomal enzymes results in sleep loss and increased oxidative stress, which are rescued through antioxidant feeding (Fig. 7). However, the exact role of oxidation in sleep induction is a topic of intense debate. While some studies suggest that increased brain oxidation induces sleep[2,5,48], those works do not provide direct evidence that the induction of brain oxidation is sleep-promoting[2] or are focused on a specific brain region[5], or map the sleep-inducing effects of oxidation to the gut[48]. Interestingly, genetic manipulations that modulate oxidation bidirectionally result in sleep reductions. For example, overexpression of glucose-6-phosphate dehydrogenase (G6PD) results in increases in mitochondrial oxidation in Pdf neurons with a dramatic sleep reduction; this behavioral effect was also obtained after loss of G6PD[49]. Together with previous research, our findings highlight that precise control of brain oxidative conditions is fundamental to maintaining normal sleep and wake cycles, and that deviation of such oxidative homeostasis results in sleep loss.

Our data also suggest that ether-lipid and plasmalogens modulate brain oxidation. Based on their structural properties, plasmalogens have been proposed to terminate lipid peroxidation[50]. This result is consistent with the fact that PBD patients show elevated levels of oxidative markers like nitric oxide (NO) and MDA[51], and high MDA levels could be normalized with plasmalogen supplementation[52]. The fact that *Gnpat* and *Pex16* knockdown increased MDA levels supports the role of peroxisomes and plasmalogens in ROS regulation during sleep. Rescue of the reduced sleep in *Pex16* knockdown with antioxidant supplementation indicates that the oxidative damage contributes to sleep loss. Another potential role of plasmalogens in sleep regulation may be reflected in the structural changes of the plasma membrane that affect circuit function. Lipid rafts are enriched in plasmalogens[50] and are known to be hubs for signal transduction processes[53]. In addition, peroxidized lipids directly regulate the potassium channel *Shaker* activity in sleep-promoting neurons[5]. And during bacterial infection, peroxisomes alter the plasma membrane lipid composition to promote the release of inflammatory cytokines[54]. Therefore, in addition to their antioxidant function, peroxisomes, through the synthesis of ether lipids and plasmalogens, may modulate the function of sleep-regulating circuits.

Although peroxisomes are ubiquitous organelles, their role in the nervous system function has been mainly explored from a glial perspective. In flies and mice, glial loss and gain of function of *Acox1* result in glial and axonal loss[55], and animals lacking the peroxisomal import protein *Pex5* in glia develop axon degeneration and neuroinflammation[56,57]. Our work shows a role of neuronal peroxisomes in modulating behavior and suggests a potential functional compartmentalization of neuronal and glial peroxisomes. Consistent with previous reports that show that *Acox1* is only expressed in glia[55], we found that *Acox1* knockdown in different neuronal subpopulations did not result in substantial sleep reduction. In addition, knockdown of *Pex16* using the pan-glial driver, repo, did not reduce sleep. The fact that peroxisomal knockdown in glia does not affect sleep (Fig. 2), but previous reports show a profound impact on neurodevelopment[12], suggests that glial peroxisomes play a more structural and metabolic role in the brain. In contrast, neuronal peroxisomes may have more signaling roles, responding and adapting their metabolism to changes in neuronal activity, oxidative state, and behavior.

Interestingly, the effect of peroxisomal knockdown on sleep is not homogeneous across the fly brain. Knockdown of *Pex16* in different neuronal subpopulations results in diverse effect sizes.

While this could reflect driver strength, it is also possible that peroxisomes are more relevant in modulating neuronal physiology in certain neuronal types, such as the mushroom body output neurons or the EB neurons. Sensitivity to metabolic changes has been reported in other sleep-promoting circuits in the fly brain. For example, the activity of the dorsal-fan-shaped body neurons is modulated by mitochondrial respiration byproducts, which are sensed by a subunit of the *Shaker* potassium channel[4,58]. Differential sensitivity to peroxisomal metabolism is also consistent with the fact that pan-neuronal *Pex16* knockdown does not result in a decrease in sleep, perhaps because the loss in other neuronal populations masks the impact of peroxisomal dysregulation in relevant brain regions.

Peroxisomal modulation of sleep can also be sex specific. For example, we found that peroxisomal numbers in females remain unchanged after sleep deprivation (Fig. S4D). This effect could reflect sex-specific differences in lipid composition. For example, VLCFAs and Ultra-Long-Chain Fatty Acids are enriched in female neurons, and their levels increase after sleep deprivation[34]. This VLCFA accumulation, together with our data, suggests that females do not rely on peroxisomal metabolism under conditions of sleep deprivation, but peroxisomes are still important in modulating baseline sleep. In addition, loss of *Pex1* and *Pex13* have sex-specific effects on lifespan and ROS accumulation[59] which highlights that in general, peroxisomal biogenesis and metabolism have a sex-specific component. Sex-specific differences in our results also reflect the sexual dimorphism in sleep amount[60]. In fact, the observed lack of sleep rescue after antioxidant supplementation in males may reflect a ceiling effect. Even though loss of *Pex16* results in an overall sleep reduction, the remaining sleep amount could be high enough to prevent further increases after antioxidant supplementation. However, our results indicate that the peroxisomal regulation of baseline sleep through control of ROS is overall conserved in both males and females.

It is intriguing that knockdown of peroxisomal genes in the ellipsoid body results in brain-wide changes in oxidation. From a circuit perspective, changes in the function of the ellipsoid body neurons could propagate to downstream neurons in the network, resulting in a systemic effect[61,62]. An alternative explanation that is consistent with neuronal-glia coupling, is that changes in neuronal activity or function due to peroxisomal dysfunction are propagated to glial cells, and the brain-wide increase in lipid peroxidation occurs mostly in glial cells because of changes in neuronal signaling. Both possibilities highlight how activity in a discrete brain region may affect the whole brain.

We found that daily oscillations in peroxisomal abundance track behavioral activity and are affected by sleep deprivation (Fig. 4). However, intrinsic clock-dependent and clock-independent rhythms can also contribute to this change in abundance. Without a functional clock, *Pex14* and *Abcd3* expression show a 12-hour rhythm in the larval fat body that follows ROS oscillations[63]. In addition, *Drosophila* functional analogs of peroxisome proliferator-activated receptors (PPARs), which include *Hnf4*, E75, and E78,[64] have been implicated in sleep and circadian rhythm regulation[65–67]. A rhythm in peroxisomal function is consistent with the temporal compartmentalization of cellular processes, where reductive processes are temporally separated from oxidative processes[68]. In animals, this separation may coincide with sleep and wake cycles. This could explain why the effect of NACA was higher at night than during the day.

In summary, we find direct evidence that sleep regulates peroxisomal abundance and metabolism, and that neuronal peroxisomes regulate baseline sleep. Among the main peroxisomal metabolic processes, we discovered that ether-lipid and plasmalogen synthesis are the most relevant sleep regulators and play a role in reducing brain oxidation. This work positions the peroxisome as a relevant component of the cellular basis of sleep.

## Materials and methods

### Drosophila melanogaster

Flies were raised on a standard cornmeal-molasses diet with the following composition: 64.7 g/L cornmeal, 27.1 g/L dry yeast, 8g/L agar, 61.6mL/L molasses, 10.2 mL/L 20% tegosept, and 2.5 mL/L propionic acid. For the thermogenetic experiments (involving TRPA1), flies were raised at 18°C; otherwise, flies were raised at 25°C. All the light conditions were 12:12 LD.

The following fly stocks were used in this study: *iso^31^*(control strain for thermogenetic experiments, lab stock), *wcs* (control strain, lab stock), Tub-PMP34 cerulean (BDSC #64246); UAS-Catalase RNAi (BDSC #31894), UAS-Acox1 RNAi (BDSC #52882), UAS-Gnpat RNAi (BDSC #52914), UAS-ABCD3 RNAi (BDSC #34349), UAS-Pex16 RNAi (BDSC #57495), UAS-GFP-SKL (BDSC #28881), R23E10-Gal4 (BDSC #49032), MB011B-Gal4 (BDSC #68294), MB051B-Gal4 (BDSC #68275), R58H05-Gal4 (BDSC #39198), c584-Gal4 (BDSC #30842), Nsyb-Gal4 (lab stock), Repo-Gal4 (lab stock), teashirt-Gal80 (Gift from Dr. Rebecca Chung-Hui Yang, FBti0114123), R23E10-GeneSwitch (lab stock), UAS-TrpA1 (Gift from Dr. Leslie Griffith, maintained as lab stock outcrossed to *iso^31^*, FBtp0040248), UAS-RedStinger (BDSC #70750), R58H05-p65.AD (BDSC #8546), R48H05-Gal4.DBD (BDSC #69353).

### Sleep and locomotor activity

Mated 5-7-day-old flies were loaded in tubes containing 5% sucrose and 2% agar. For the NACA, gaboxadol and GeneSwitch experiments, the food was supplemented with 160 μg/mL of NACA (Sigma-Aldrich A0737), 0.1 mg/mL Gaboxadol (Sigma Aldrich, T101), and 0.2 mg/mL Mifepristone (Sigma Aldrich, M8046) respectively. Sleep was recorded using the Drosophila Activity Monitoring (DAM) system (TriKinetics, Waltham, MA). The single-beam version (DAM2) was used for the experiments involving pharmacological manipulations and thermogenetic activation. Otherwise, the flies were loaded on multi-beams monitors (DAM5H). All flies were recorded in a LD 12:12 light schedule at 25°C except for the TRPA1 experiments, where flies were recorded at 31°C during the activation day. Sleep was analyzed using custom Matlab[69] scripts, and sleep bouts were defined as 5 continuous minutes of inactivity. For the analysis, the first two recording days were discarded, and the mean of the three following days was calculated for the comparisons between the groups. For experiments involving TRPA1 activation, sleep was calculated as the ratio between sleep during activation and baseline sleep, with baseline sleep being the total sleep time of the first day of recording at 25 °C. We defined that our manipulation had effects on sleep if and only if the experimental group was significantly different from both the Gal4 and the UAS control groups. The screening plots were made by subtracting the mean of each experimental control from the mean of its corresponding Gal4 and UAS control.

### Brain dissections and immunohistochemistry

For endogenous fluorescence signal recordings (PMP34, GFP-SKL, mcd8GFP, RedStinger), brains from mated 5–7-day-old flies were dissected in ice-cold PBS and fixed with 4% paraformaldehyde in PBS for 30 minutes at room temperature. Followed by three washes with 0.4% PBS Triton X-100 (PBST) and one quick wash with PBS. The brains were mounted on glass slides with VECTASHIELD (Vector Labs, H-1000).

For the MDA staining, after the third wash with PBST, the brains were blocked using 10% NGS in PBST for one hour at 25°C and incubated overnight at 4°C with the primary antibody (Monoclonal Anti -Malondialdehyde antibody produced in mouse. Clone 11E3, Sigma Aldrich, SAB5202544, 1:100). One group was incubated without primary antibody to use as a negative staining control. After incubation, brains were washed 3 times with PBST and incubated with the secondary antibody (Goat anti-Mouse IgG (H+L) Cross-Adsorbed Secondary Antibody, Alexa Fluor^TM^ 594, ThermoFisher Scientific,1:250) for 2 hours at 25 °C and protected from light. After the secondary antibody incubation, the brains were washed three times with PBST, followed by a quick wash in PBS. The brains were mounted on glass slides with VECTASHIELD.

For the Bodipy^TM^ 581/591 C11 experiments, brains were dissected in ice-cold complete media (Schneider’s Drosophila Medium + Fetal Bovine Serum 10%), washed once with complete media, and incubated with 2μM Bodipy^TM^ 581/591 C11(Invitrogen, D3861), in complete media for 30 minutes at 25°C. After incubation, brains were washed three times with complete media and mounted on glass slides with VECTASHIELD for immediate imaging.

All the preparations were imaged using a Leica Stellaris STED confocal microscope with a 40X oil objective, and z-stacks spanning the whole brain or the ellipsoid body (for the ring GFP-SKL imaging) were acquired.

### Sleep deprivation experiments

Flies in vials with complete food were loaded in a Vortexer Mounting Plate (TriKinetics-TVOR 120) and mechanically sleep deprived through application of random 2-second pulses every 20 seconds. The intensity was adjusted so that during the deprivation pulses, the flies were sent to the bottom of the vial but were able to climb to the walls during the interpulse interval. Sleep deprivation was made in 6-hour windows (ZT8-ZT14 and ZT14-ZT20).

### Lipidomic data analysis

Neuron and glia-specific lipidomic data were obtained from previous work[34]. From that dataset, phospholipids that changed abundance with wake (ZT14) or after sleep deprivation (ZT2 SD) relative to ZT2 with an FDR < 0.01 were selected. To classify a phospholipid as potentially peroxisomal-derived, a manual inspection of the predicted lipid species was performed. If the predicted species contains a predicted ether or vinyl ether bond, it was classified as an ether lipid or plasmalogen, respectively, and included in the peroxisomal-derived category. For those instances where non-peroxisomal-related phospholipids were also detected, the percentage of each category was calculated.

### NACA treatment

N-Acetylcysteine amide was diluted in milliQ water to a stock concentration of 10mg/mL and diluted to a final concentration of 160μg/mL in complete food or in 5% sucrose, 2% agar food. Flies were pre-exposed to NACA in complete food in vials for three days and then loaded in vials with sucrose/agar food with NACA for the sleep recording. For the imaging experiments involving NACA treatment, flies were only exposed to NACA in complete food. For the control groups, instead of NACA, an equivalent volume of water was added to the food.

### Drosophila brain single-cell dataset analysis

Single-cell Transcriptome dataset[70] was accessed from https://scope.aertslab.org, analysis was done from the Adult Brain Filtered 57k version, and was visualized using the Seurat 82PC, 30 perplexity coordinates. The gene *CG6024*, which is the gene promoter used to create the R58H05-Gal4 line, and *Gad1* were used as ellipsoid body identifiers. While *Pex16* and *Pmp70* were used as peroxisomal identifiers. Each ellipsoid body identifier was queried with each peroxisomal identifier to evaluate the degree of co-expression within the same cluster.

### Confocal imaging analysis

All images were analyzed using custom ImageJ macros, and the representative images per experiment were edited in a way that all have the same intensity ranges. For the PMP34 puncta counting, a maximum intensity projection of the central brain was made, and the background was subtracted using the Convoluted Background Subtraction tool from the BioVoxxel Toolbox[71]. From the resulting image, an auto threshold with the Maximum Entropy method was performed, and the Analyze Particles tool was applied to the thresholded image to count puncta. The number of puncta was normalized by ZT2 for the time-course experiments, or ZT14 and ZT20 for the sleep deprivation experiments.

For GFP-SKL quantification. A maximum intensity projection of the ellipsoid body ring was made. Background intensity was measured from a rectangular ROI in a region without fluorescence. That background intensity was subtracted from the original image projection, and the Raw Integrated Density was measured and compared between groups.

For MDA antibody signal quantification. A sum of slices of the central brain was made, and the mean gray value was measured. When the number of slices in the stack was not equal, the mean gray value was normalized by the number of stacks. To account for background intensity, the average slice-normalized mean gray value from the negative staining control was subtracted from all the groups. Data was normalized with the Gal4 control’s signal. For the representative images, the Fire LUT was applied.

All the macros for analysis are deposited in: https://github.com/CamiloGuevaraEsp/Imaging-quantification-scripts

### Statistical Analysis

All significance tests were performed using GraphPad Prism 10. Before analysis, the Shapiro-Wilk normality test and Bartlett’s test were performed to evaluate normality and equality of variances. If both conditions were met, in those cases where more than two groups were involved, an Ordinary one-way ANOVA followed by a Bonferroni multiple comparison correction was performed. If only the normality but not the equality of variances was met, a Brown-Forsythe ANOVA test with Dunnett’s multiple comparison correction was performed. If neither of the two conditions was met, a Kruskal-Wallis test followed by a Dunn’s multiple comparisons test was performed. If the comparison involved only two groups with a normal distribution, an unpaired t-test was performed; otherwise, the comparisons were made using a Mann-Whitney test. Significant thresholds were p-values <0.05. All the sleep experiments were repeated at least three independent times, and all the imaging experiments were repeated at least two times.

## Acknowledgments

We thank all members of the Sehgal laboratory for their valuable comments. Stocks obtained from the Bloomington Drosophila Stock Center (NIH P400D018537) were used in this study.

## Supporting information

**S1 Fig:**
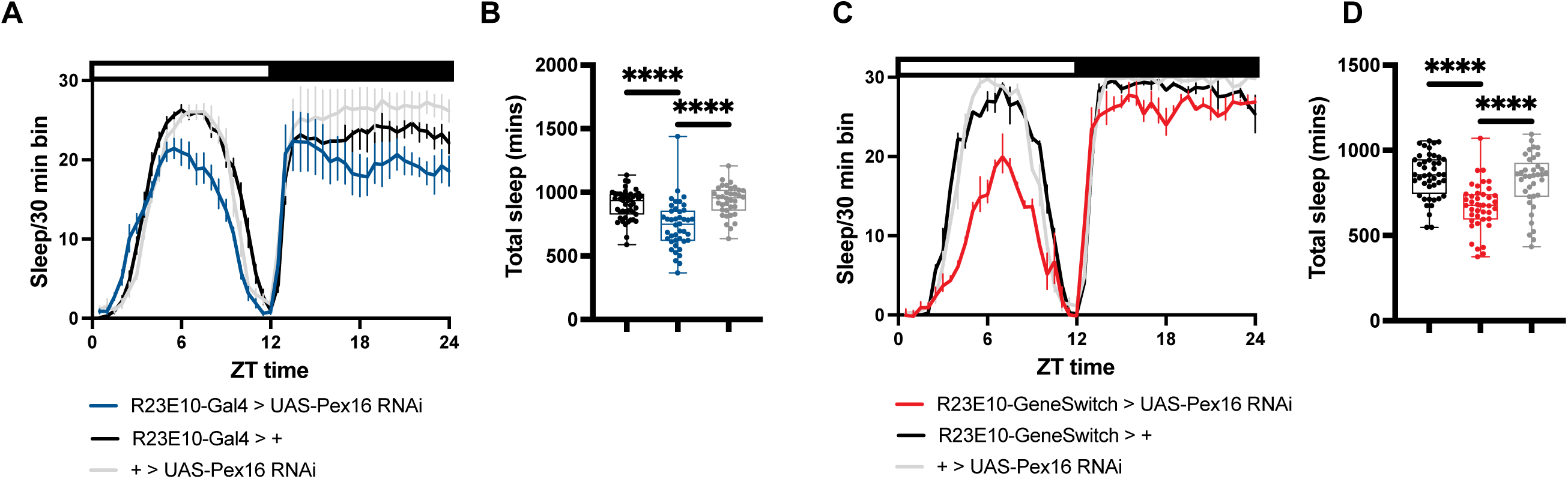
Developmental and adult-specific knockdown of *Pex16* with the R23E10 driver reduces sleep in females. **(A, C)** Female sleep trace after developmental (A) and adult-specific (C) knockdown of *Pex16* using the R23E10-Gal4 and R23E10-GeneSwitch drivers, respectively. **(B, D)** Total sleep after developmental (B) (n= 36-47) and adult-specific (D) (n= 37-44) knockdown of *Pex16* using the R23E10-Gal4 and R23E10-GeneSwitch drivers, respectively. For figure B, \*\*\*\**p* < 0.001 by Kruskal-Wallis test with Dunn’s multiple comparisons test. For figure D, \*\*\*\**p* < 0.001 by Ordinary one-way ANOVA with Bonferroni multiple comparisons correction.

**S2 Fig:**
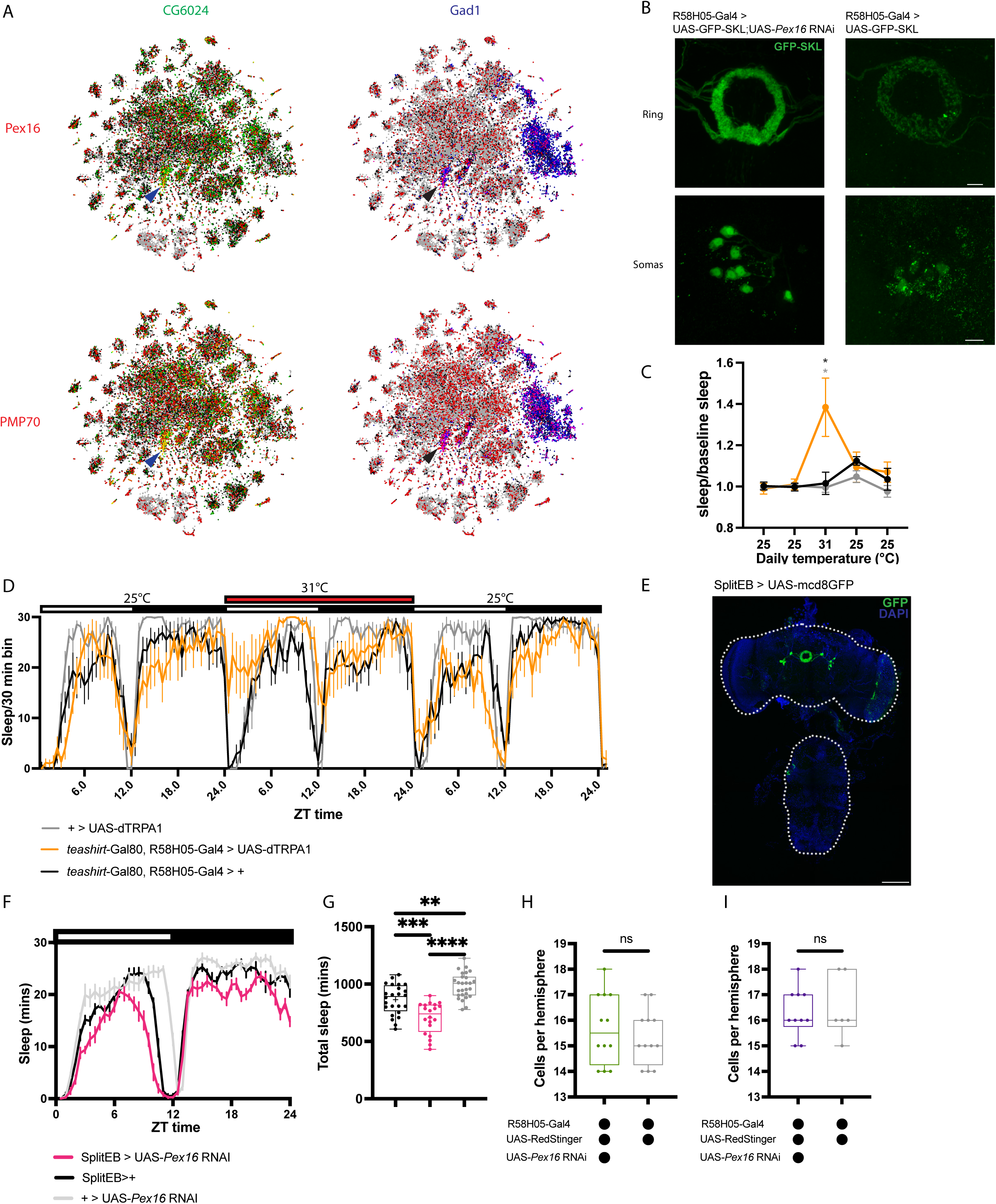
Validation of the ellipsoid body as a region to study peroxisomal function and sleep. **(A)** Brain Single-cell Transcriptomic atlas visualized using the Seurat 82PC, 30 perplexity coordinates. Blue arrowhead points to the cluster with colocalization between *CG6024* and *Pex16* (upper left), and *Pmp70* (bottom left). Black arrowhead points to the cluster with colocalization between *Gad1* and *Pex16* (upper right), and *Pmp70* (bottom right). **(B)** Peroxisomal GFP imaging in the ellipsoid body ring neuropil and somas with (left) and without (right) *Pex16* knockdown. Scale bar for ring neuropil and cell bodies is 10μm. **(C)** Total sleep time relative to baseline sleep for brain-specific activation of the R58H05 driver (n= 7-8), *p < 0.05 by Ordinary one-way ANOVA with Bonferroni multiple comparisons correction. **(D)** Sleep trace for the experiment mentioned in (C). **(E)** Expression pattern for the R58H05-AD; R48H04-DBD split driver line. Scale bar is 100μm. **(F)** Male sleep trace for *Pex16* knockdown in the split driver line. **(G)** Total sleep time after *Pex16* knockdown in the split driver line (n= 20-27), **p < 0.01, ***p < 0.001, ****p < 0.0001 by Ordinary one-way ANOVA with Bonferroni multiple comparisons correction. **(H, I)** Total cell counts with and without *Pex16* loss in the R58H05 driver in males (H) (n = 12) and females (I) (n= 6-10). Unpaired t-test.

**S3 Fig:**
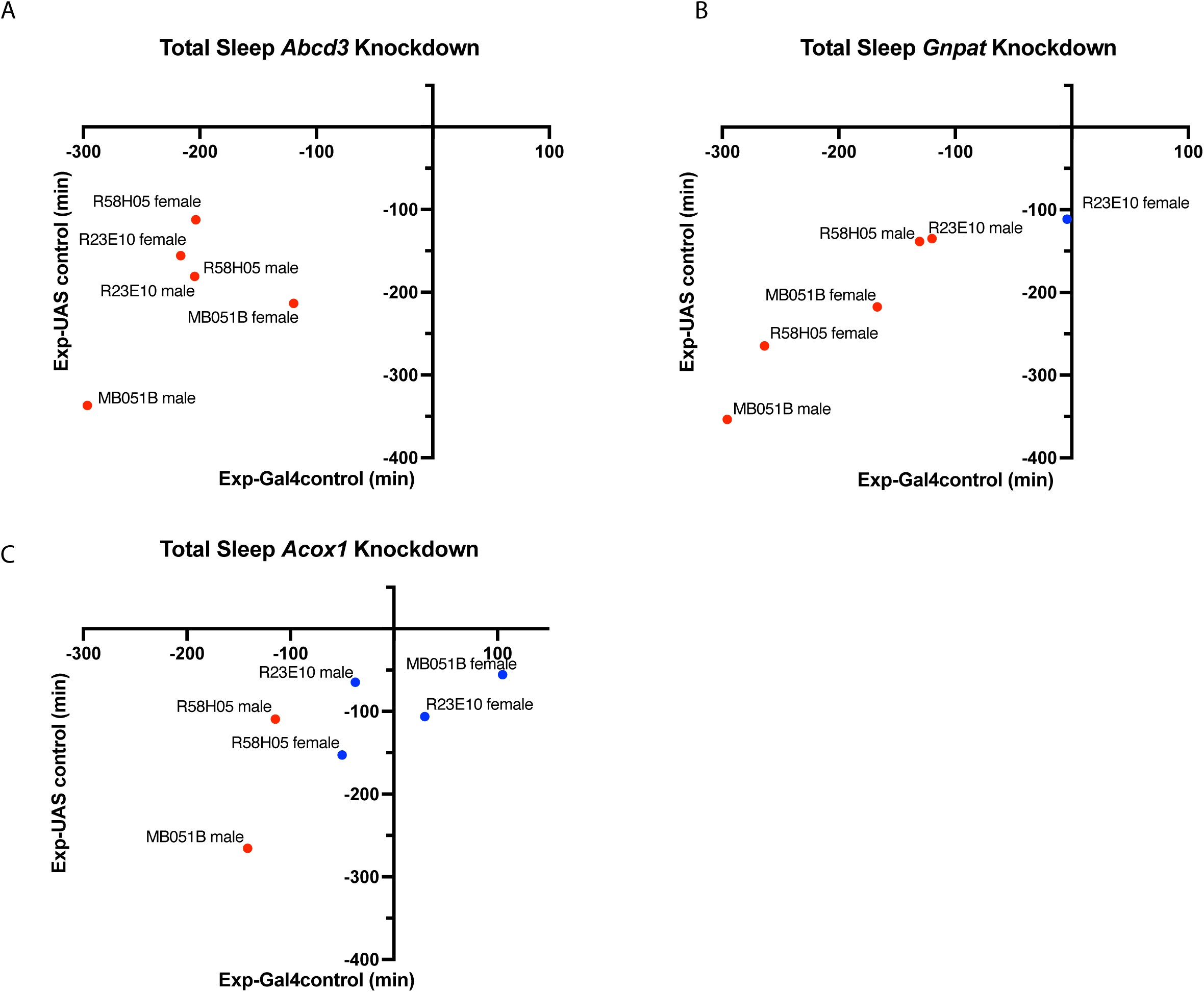
Knockdown of peroxisomal lipid import and synthesis, but not beta-oxidation, in sleep-promoting regions reduces sleep. Sleep differences after knockdown of *Abcd3* **(A)**, *Gnpat* **(B)**, and *Acox1* **(C)** in different sleep-promoting brain cell populations in males and females. X and Y axes represent the sleep differences between the knockdown group and the Gal4 and UAS control, respectively. Red dots indicate significant sleep time differences compared with the UAS and Gal4 controls (n=41-48), while blue dots represent non-significant differences. R23E10: dorsal fan-shaped body and ascending neurons driver; R58H05: ellipsoid body driver; MB051B: Sleep-promoting mushroom body output neurons driver.

**S4 Fig:**
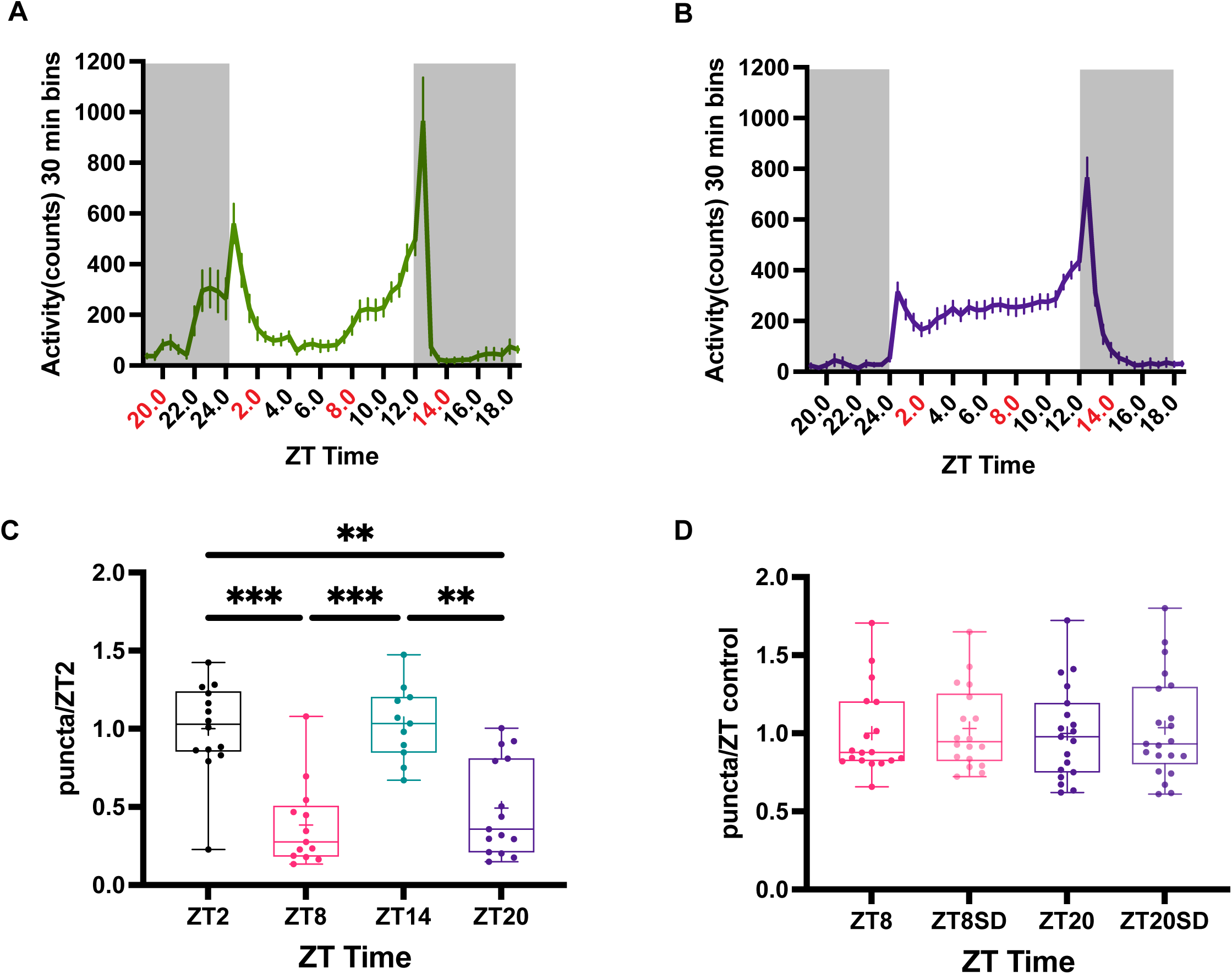
Timepoints with increased peroxisome abundance correspond to periods of higher behavioral activity, but sleep deprivation does not affect peroxisomal abundance in female flies. **(A,B)** PMP34-mCerulean male (A) and female (B) activity traces. Red numbers show the time points where peroxisomal abundance was measured. **(C)** Quantification of mCerulean puncta in female flies relative to ZT2 (n= 11-15), \*\**p* < 0.01, \*\*\**p* < 0.001 by Kruskal-Wallis test with Dunn’s multiple comparisons test. **(D)** Quantification of mCerulean puncta relative to their corresponding non-sleep-deprived control (n= 18-21). Mann-Whitney test between ZT8 and ZT8SD, unpaired t-test between ZT20 and ZT20SD.

**S5 Fig:**
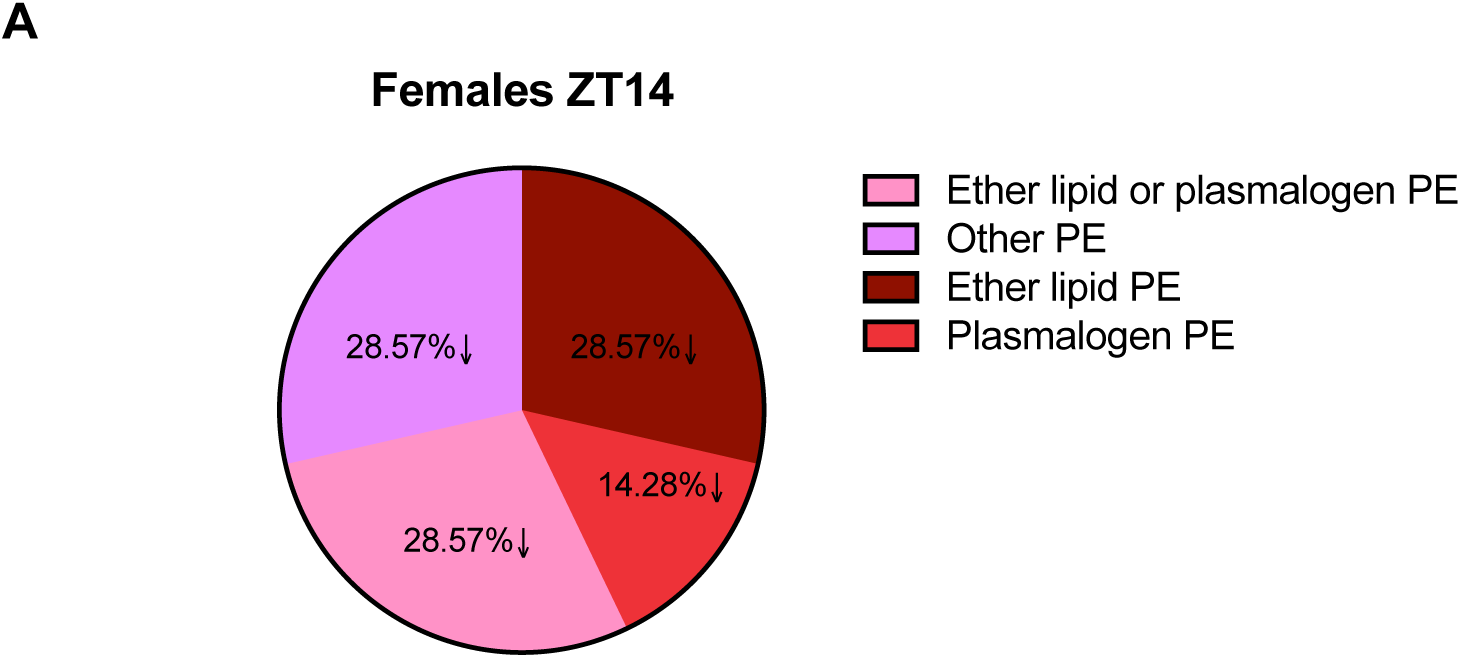
In glia, peroxisome-derived phospholipids are only detected in females after wake. **(A)** Percentage of peroxisomal-related phosphatidylethanolamines (PE) showing a significant change in abundance compared to ZT2 in glia. The arrow shows a decreased abundance relative to ZT2.

**S6 Fig:**
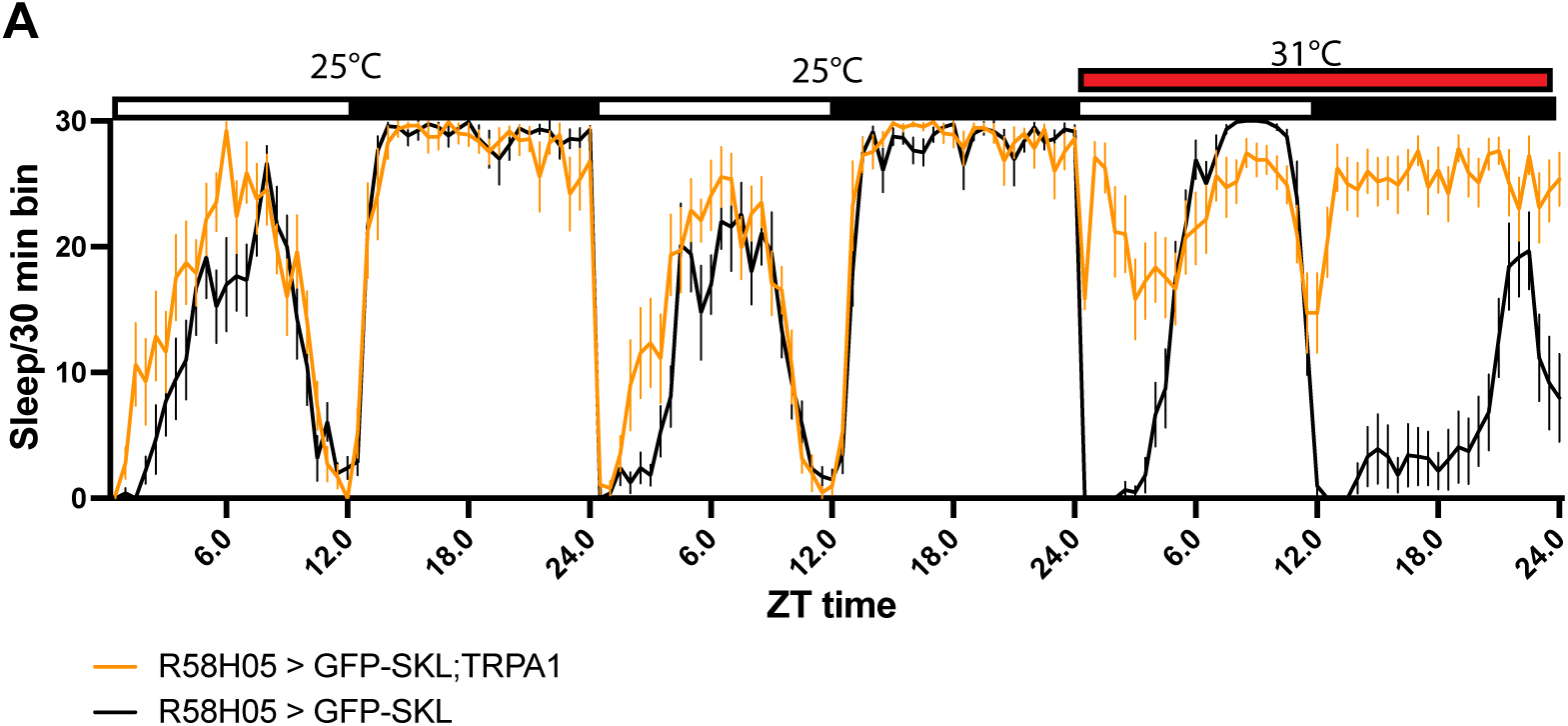
Validation of neuronal activation in GFP-SKL flies. **(A)** Female sleep trace of GFP-SKL expressing flies with and without neuronal activation through TRPA1. Flies for figures 5A, 5E were dissected immediately after the end of the 31 °C pulse.

**S7 Fig:**
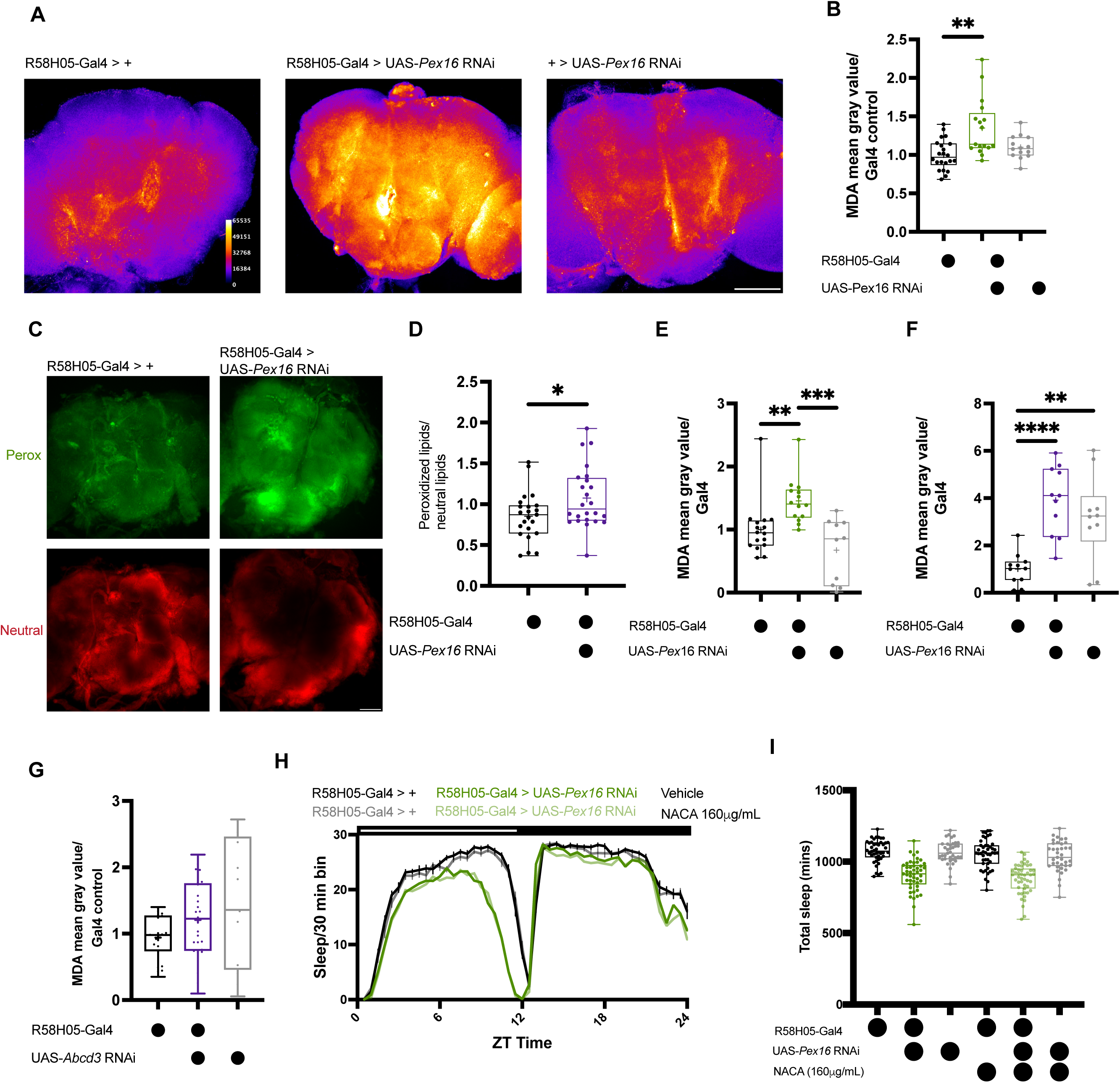
Increase in brain oxidation after knockdown of peroxisomal genes is independent of sleep loss and lipid import into the peroxisome. **(A)** Representative images of Bodipy^TM^ 581/591 C11-stained brains with (right column) and without (left column) *Pex16* knockdown in the R58H05 driver. The upper row shows the peroxidized lipids signal, and the lower row shows the neutral lipids signal. Scale bar is 100 μm. **(B)** Quantification of the ratio between peroxidized and neutral lipids in the conditions described in (A) (n= 23-24), *p < 0.05 by Unpaired t-test. **(C)** Quantification of the mean gray value normalized to the Gal4 control after *Pex16* knockdown in the R58H05 driver with Gaboxadol 0.1mg/mL supplementation (n= 10-16), in males **p < 0.01, ***p < 0.001 by Kruskal-Wallis test with Dunn’s multiple comparisons test. **(D)** Quantification of the mean gray value normalized to the Gal4 control after *Pex16* knockdown in the R58H05 driver with Gaboxadol 0.1mg/mL supplementation (n= 10-11), in females **p < 0.01, ****p < 0.0001 by One-way ANOVA with Bonferroni multiple comparisons correction. **(E)** Quantification of the mean gray value normalized to the Gal4 control after *Abcd3* knockdown in the R58H05 driver in females (n= 13-25), Brown-Forsythe ANOVA test with Dunnett’s multiple comparison correction.

## Notes

### Competing Interest Statement

The authors have declared no competing interest.

